# Predicting the re-distribution of antibiotic molecules caused by inter-species interactions in microbial communities

**DOI:** 10.1101/2020.12.14.422780

**Authors:** Carlos Reding

## Abstract

Microbes associate in nature forming complex communities, but they are often studied in purified form. Here I show that neighbouring species enforce the re-distribution of carbon and antimicrobial molecules, predictably changing drug efficacy with respect to standard laboratory assays. A simple mathematical model, validated experimentally using pairwise competition assays, suggests that differences in drug sensitivity between the competing species causes the re-distribution of drug molecules without affecting carbon uptake. The re-distribution of drug is even when species have similar drug sensitivity, reducing drug efficacy. But when their sensitivities differ the re-distribution is uneven: The most sensitive species accumulates more drug molecules, increasing efficacy against it. Drug efficacy tests relying on samples with multiple species are considered unreliable and unpredictable, but study demonstrates that efficacy in these cases can be qualitatively predicted. It also suggests that living in communities can be beneficial even when all species compete for a single carbon source, as the relationship between cell density and drug required to inhibit their growth may be more complex than previously thought.

## I. Introduction

The notion of pure culture is fundamental in microbiology. The isolation and growth of individual species is justified by the seemingly unpredictability (*1*) of *in vitro* assays when cultures contain multiple species. For antimicrobial sensitivity tests, this means the same microbe can show different sensitivities to the same drugs, depending on whether the cultures contain one or multiple species (*2-4*). But microbes live in communities in nature, even when they are mostly competing for resources (*5*). The question is, therefore, whether co-existing species can threaten drug efficacy *in vivo* with respect to standard sensitivity tests. Current data suggest they do (*4, 6-9*), notably reducing drug efficacy, and the result is that infections containing multiple species are more difficult to treat (*10*−*12*)—requiring alternatives to antibiotics altogether (*10, 12*)—and waters more difficult to treat (*13*). This phenomenon has been associated to biofilm formation (*14, 15*), stochastic phenotypic variations of isogenic bacterial populations (*14, 15*), signalling molecules (*14, 15*) or enzymatic degradation of antimicrobials (*14, 15*). But questions still remain about its predictability, and why, sometimes, drug efficacy seems to increase (*6*).

Below I present a model that provides a passive physical mechanism to explain and, perhaps more importantly, predict this phenomenon. The model relies on Fick’s first diffusion law and aims to understand how the microbial growth changes the flow of carbon and antimicrobial molecules, and, thus, influence drug efficacy against all species exposed. Note that, when surrounded by neighbours, species attain lower densities within a community with respect to that in pure culture because carbon is shared between multiple species (*16*). Now, the model predicts that carbon and antimicrobial molecules will distribute evenly among all species exposed if they have similar sensitivity to the antimicrobial. The result is relatively less drug molecules per cell of each species, and therefore lower drug efficacy. If one or more species are *not* sensitive to the antimicrobial, carbon molecules still distribute evenly as it is an active process (*17, 18*) but drug molecules are not: They flow back through diffusion from non-sensitive species into the environment, re-exposing those that are sensitive to the drug. Here, the model predicts relatively more drug per cell of sensitive species resulting thus in higher drug efficacy. These predictions were maintained across a range of parameter values for carbon uptake rate, carbon affinity, and biomass yield—fundamental components of the growth function in microbes (*19*).

To validate these predictions, I measured the efficacy of tetracycline against *Escherichia coli* Wyl in standard sensitivity assays using pure cultures, cultures with equal proportion of another microbe with similar drug-sensitivity (*Salmonella typhimurium*), and cultures containing equal proportion of another microbe now tolerant to tetracycline (*Escherichia coli* GB(c)). Consistently with the theoretical predictions, inhibiting Wyl required more tetracycline—with respect to pure cultures—in the presence of equally sensitive neighbours and less tetracycline in the presence of drug-tolerant neighbours.

## II. Results

Consider *n* phenotypically distinct species competing for a limited resource, *C*, and exposed to a drug, *A*, supplied at concentration *A*_*e*_ (0) = *A*_0_, cast as the following model:

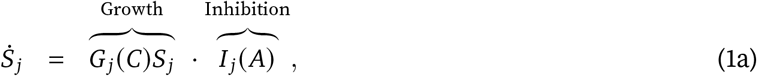

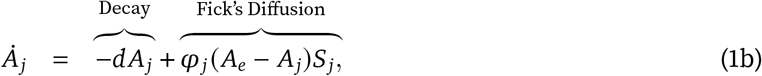

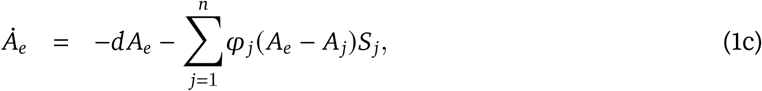

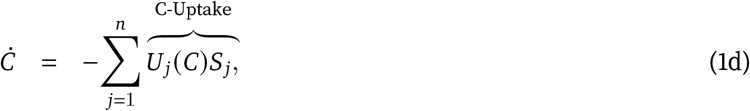

Here, 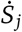 and 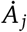 represent the density of individuals per unit volume from species *j* and their content of drug *A* over time, respectively, with initial conditions *S*_*j*_ (0) = *S*_*j*0_, *A*_*j*_ (0) = 0, and *C*(0) = *C*_0_ > 0. The uptake of resource *C* by individuals of species *j* can be modelled as a saturating Monod function given that carbon transport is mediated by enzymes in nature (*17, 18*). This function is proportional to the maximal uptake rate 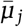,

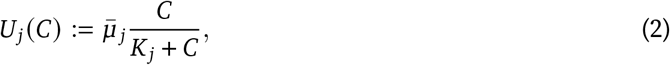

where *K*_*j*_ is the half-saturation parameter—the affinity of individuals from species *j* for the limited resource *C* is therefore given by 1*/K*_*j*_. The growth rate of each species, at a given resource con-centration is denoted by *G*_*j*_ (*C*) := *U*_*j*_(*C*) · *y*_*j*_, where *y*_*j*_ is the biomass yield per unit of resource in individuals from species *j*. Note that, for simplicity, I assume that *y*_*j*_ does not vary between pure and mixed culture conditions. Biomass yield depends on resource availability (*19*) and competition for a common carbon source will likely change *y*_*j*_ in all competing species, albeit these changes can be difficult to measure (*20*). Their growth inhibition by drug *A* is described qualitatively by the Hill function (*21*)

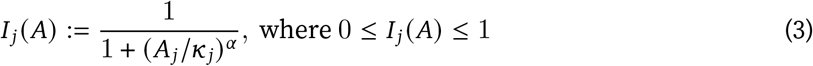

scaling the growth function *G*_*j*_ (*S*) similarly to bacteriostatic drugs (*22*). This function is dimensionless and has two parameters. First, *α* is a dimensionless Hill coefficient which characterises the co-operativity of the inhibition. And second, *κ*_*j*_ is the affinity of drug *A* for its target. It can be derived from the drug concentration required to halve the maximal growth rate, so that *A*_50_ = 1/*κ*_*j*_ → *κ*_*j*_ = 1*/A*_50_ (*21*). For the sake of simplicity, I assumed that drug *A* diffuses from the environment into cells of species *j*, and *vice versa*, following Fick’s first diffusion law (*23*) with a diffusion coefficient *φ*_*j*_; and part of *A* being lost to chemical stability (*24*) at a rate *d*. This is not an unreasonable assumption given that antimicrobials molecules can indeed diffuse through cell membranes facilitated by membrane proteins (*25*). Consequently, drug *A* is released in active form following the death of target and non-target species.

For my first computation I set the number of species *j* = 2, to facilitate later experimental validation, where *I*_1_ (*A*) = *I*_2_ (*A*) and *G*_1_ (*C*) = *G*_2_ (*C*). Thus, individuals from both species are sensitive to *A* and phenotypically identical. Given Equation 3, the density of individuals from either species as pure cultures, after 24h of incubation, declines with higher drug concentrations consistently with standard clinical protocols (*26*) (Figure 1A). To allow experimental validation, I calculated the concentration of *A* inhibiting the growth of the pure cultures by 90% (IC_90_) as commonly used in clinic laboratories (*27*−*29*). The drug sensitivity of each species depends on the values for the parameters 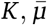, and *y* of Equation 2 (Figure 1B-D, grey), with values that increase the density of individuals resulting in higher IC_90_. This is consistent with the *inoculum effect* (*30*) (Figure S1), whereby sensitivity tests that use larger inocula also report higher minimum inhibitory concentrations, hence the standardisation of these clinical assays.

**Figure 1.**
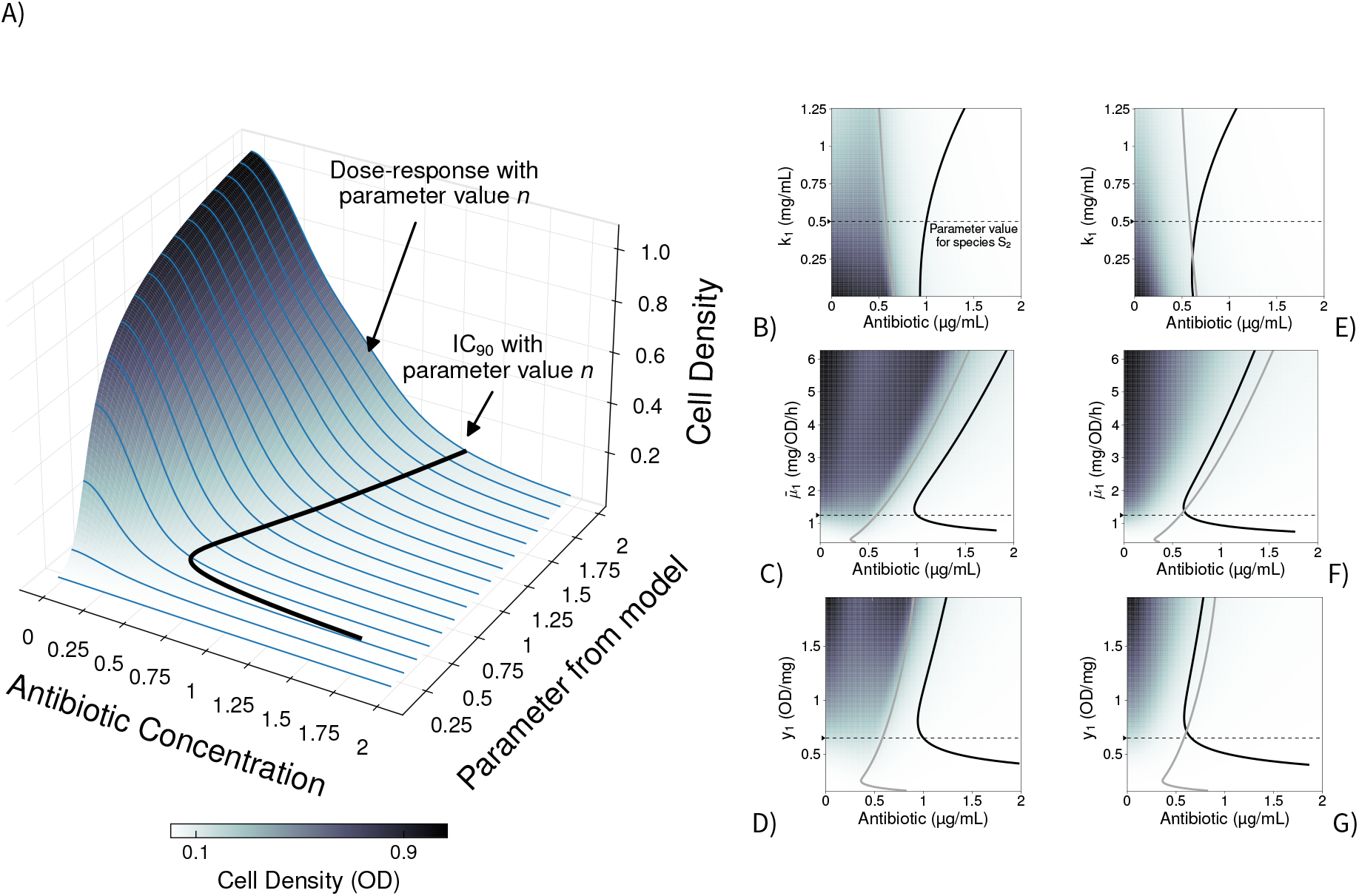
Si drug sensitivity profiles in pure and mixed culture growth conditions alongside species S2. **A)** Growth of species S1, with different parameter values (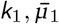, and *y*_1_), after 24h of growth in the presence of different antibiotic concentrations. I aggregated the resulting dose-response profiles (blue) to create a density map from low predicted cell density (white) to high predicted cell density (black). **B-D**)IC_90_, antibiotic concentration inhibiting 90% (IC_90_) the growth predicted without drug, resulting with different parameters values for the half-saturation parameter *k*_1_ (B), maximal carbon up-take 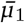 (C), or biomass yield *y*_*s*_ (D) in equation 1 when species S_2_ is drug-sensitive. The IC_90_ for species S_1_ growing as pure cultures is shown in grey, and growing in mixed culture with S_2_ are shown in black. The parameter values for species S_2_ were fixed at a value noted by a black arrow on the *y*-axis, followed by a dotted black line. **E-G)** Change in IC_90_, as in Figures B-C), when the competing species S_2_ is *not* drug-sensitive (resistant or tolerant).

Figure S1 shows explicitly the theoretical relationship between IC_90_ and cell density of species S_1_ at the IC_90_ in pure culture, direct and non-linear consistently with prior data (*30*−*32*). For parameters 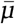 and *y*, for example, the resulting change in IC_90_ with respect to the parameter values is nonmonotone, which means certain values for these parameters can maximise susceptibility for drug *A*. The neighbouring species shows similar, albeit not identical, changes in sensitivity (Figure S2). This non-linearity is caused by cultures not reaching the equilibrium, for all drug concentrations and parameter values tested, within the standard 24h incubation times. If, in the model, cultures are allowed to grow for longer and reach the equilibrium, the IC_90_-cell density profile can change (Figure S1B). This phenomenon is exacerbated if both species grow in mixed culture conditions, where both become phenotypically more tolerant to drug *A* (Figure 1B-D, black). If I were to target, say, individuals from species *S*_1_, doing so when the species is surrounded by *S*_*2*_ would require more drug. This is the case, for example, of pancreatic ductal adenocarcinoma treated with gemcitabine when bacteria grow within the tumour’s microenvironment (*33*). More generally, genotypes analog to *S*_1_ should increase their drug tolerance when they are surrounded by similarly sensitive species. Despite the freedom in parameter values in the model, this prediction is robust to changes in biomass yield (*y*_*i*_) or maximal carbon uptake 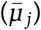 of species *S*_2_ (Figure S3).

To test this hypothesis, I mixed equal proportions (cell/cell) of *Escherichia coli* Wyl and *Salmonella typhimurium* SL1344 in minimal media supplemented with different concentrations of tetracycline (see Methods). Note that the inoculum effect is not linear with respect to the size of inocula (*31, 32*) unless the difference between inocula is 100- to 1000-fold or more, which is not the case here. Tetracycine can diffuse passively into cells of both gram-negative species (*34*), who also have similar sensitivity to this antibiotic: 0.232±0.003 and 0.276±0.016 μg/mL of tetracycline (mean IC_90_ ±95% confidence, with *n* = 8 replicates, see Figures S2 and S7). This approximates to *I*_1_ (*A*) ≈ *I*_2_ (*A*), as laid out by the theory above. The chromosome of *E. coil* Wyl carries *yfp*, gene encoding a <u>y</u>ellow <u>f</u>luorescence <u>p</u>rotein (YFP), so I tracked its density in mixed culture conditions. Consistently with equations 1a-d, the bacterium was around 23% more tolerant to tetracycline when it grew in mixed culture with *S. typhimurium* (Mann Whitnay U-test *p* = 1.554 × 10^−4^, *ranksum* = 36 with *n* = 8 replicates, Figure 2A).

**Figure 2.**
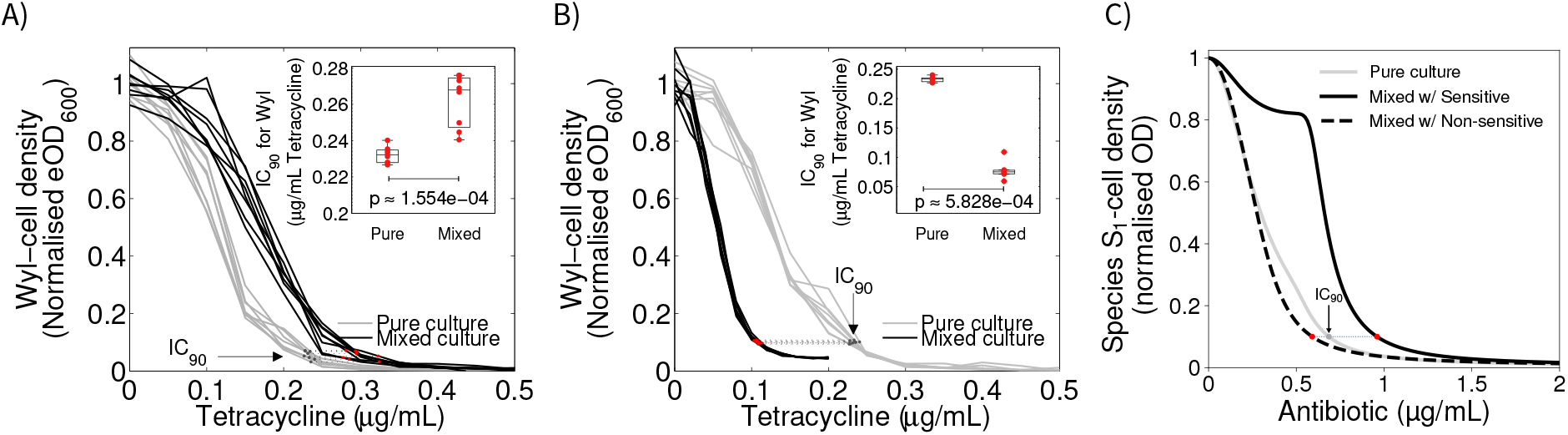
Change in drug efficacy against sensitive *Escherichia coli* Wyl are consistent with theoretical predictions. **A–B)** Change in normalised density of *Escherichia coli* Wyl as a function of tetracycline concentration, when Wyl grows in mixed culture with tetracycline-sensitive *Salmonella typhimurium* (A) and tetracycline-resistant *Escherichia coli* GB(c) (B). The change in density of Wyl growing in mixed culture is shown in black, with grey showing the change in density in pure culture. The IC_90_ in each condition is shown as dots, red for mixed culture conditions and dark grey for pure culture, connected by a dotted line. Non-parametric, Mann-Whitney U-test between IC_90_s is shown in the inset. Raw data is shown as dots, whereas the boxes represent median (centre of the box), 25th, and 75th percentile of the dataset. The whiskers show the most extreme data points that are not outliers. **C)** Theoretical change in S_1_-cell density with increasing antibiotic concentration in pure (grey) and mixed (black) culture conditions with neighbours that have different drug sensitivity. The plot represents the case where both species have different carbon uptake 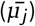, and differences in IC_90_ are represented as shown in A–B for consistency.

Next, I explored in the model the case where individuals from both species have different sensitivities to drug *A* (*I*_1_(*A*) ≠ *I*_2_(*A*)). This scenario is akin to pathogens such as *C. difficile* growing alongside human cells (*35*) where the latter are unaffected by the drug (*I*_2_(*A*) ≈ 1). The model now predicts a subset of values for *K, y*, and 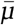 that make S_1_ *more* sensitive to the drug in the presence of individuals from species *S*_2_ (Figure 1E–G), whether *S*_2_ is not affected by drug *A* or is resistant to it for example through efflux pumps (*36*) (see Supplementary Text). To test this prediction, I mixed equal proportions (cell/cell) of two constructs of *Escherichia coli* with different sensitivities to tetracycline. One construct is Wyl, used above, who is sensitive to the antibiotic. The other construct is GB(c), harbouring a non-transmissible plasmid carrying the gene *tet* (*36*) (*37*) and, therefore, resistant to the drug. Tetracycline binds to the bacterial ribosome, inhibiting protein synthesis (*38*), and *tet*(*36*) provides ribosomal protection against tetracycline (*37*) without degrading the antibiotic. The IC_90_ for this construct was 6.106 ± 0.272 μg/mL of tetracycline (mean IC_90_ ± 95% confidence with *n* = 8 replicates). Now, *I*_1_(*A*) ≪ *I*_2_ (*A*) satisfies the assumption above. The IC_90_ for *E. coli* Wyl was 0.232 ± 0.003 μg/mL of tetracycline as pure culture. Growing alongside drug-resistant GB(c), however, it was 0.112 ± 0.003 μg/mL (Figure 2B).

It is noteworthy to highlight the two processes at play. On the one hand, the growth of each species is limited due to competition for the same resource—also known as competitive suppression (*16*). It could be conjectured that the observed variations in IC_90_ are caused by variations in growth, with respect to pure culture conditions, caused by competitive suppression. Indeed, *E. coli* Wyl grows less, with respect to pure culture conditions, in the presence of either competitor as Figure S4 illustrates. Within this difference, Wyl grows more in the presence of tetracycline-resistant *E. coli* GB(c), and yet, drug efficacy against Wyl was highest in the presence of this competitor. Now, the relative abundance of each species during the competition determines the flow and distribution of antimicrobial molecules given they diffuse passively following Fick’s law. Since glucose is actively transported into the cytoplasm, even against concentration gradients (*17, 18*), the model predicts virtually no effect on the carbon uptaken by each species as Figure S5 illustrates. In other words, the carbon source *C* is depleted more rapidly when more species use it, limiting the growth of each competing species with respect to pure culture conditions.

On the other hand, the sensitivity to drug *A* determines the fraction of carbon that gets used by each species. When *I*_1_ (*A*) *≈ I*_2_(*A*), equations 1a–d suggest that individuals from both species accumulate similar amounts of drug *A* (Figure S6A–C). This means that, analog to the carbon, drug *A* is depleted more rapidly limiting the exposure of the target species S_1_. When combining both processes for carbon and drug molecules, the result is relatively fewer of drug molecules per cell in both species with respect to that predicted in pure culture. However, when *I*_1_ (*A*) ≠ *I*_2_ (*A*), by virtue of different affinities of drug *A* for each species (*κ*_1_ ≠ *κ*_2_), the accumulation of drug molecules is uneven (Figures S6D–I). Soon after the exposure to drug *A*, the species S_2_, with the least affinity for it, and the environment are in equilibrium. In other words, the number of drug molecules that diffuse into S_2_-cells are the same that diffuse from these cells back to the environment. These molecules, in turns, are accumulated by the species with the most affinity for the drug. Here, integrating both processes results in more molecules per cell in the species with highest drug-sensitivity (S_1_).

To verify this hypothesis, I estimated the content of tetracycline in *E. coli* Wyl by dividing the bacterium’s culture density, measured in relative fluorescence units to allow tracking in mixed culture conditions, by the concentration of tetracycline defining its IC_90_. The estimates resemble closely the theoretical predictions in Figure 2C: *E. coli* Wyl contains approximately 20% less tetracycline growing next to *Salmonella typhimurium* and 65% more tetracycline growing alongside drug resistant GB(c) (Figure 3).

**Figure 3.**
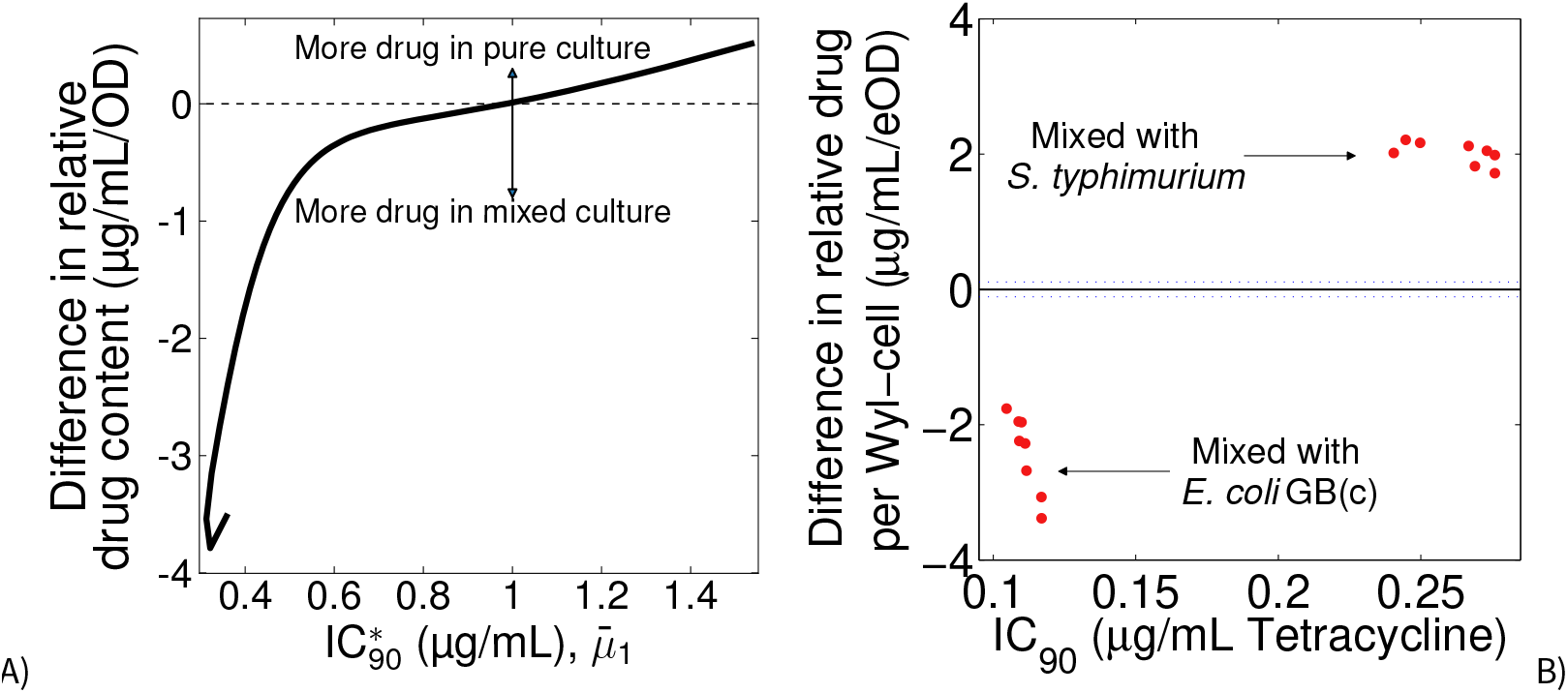
Relative content in drug molecules changes as predicted by model. **A)** Difference in drug content per S_1_-cell at the IC_90_ between pure culture and mixed culture conditions. Positive and negative values denote more drug in S_1_-cells in pure and mixed culture conditions, respectively. Lack of difference is shown as a horizontal, dotted line. **B)** Experimental estimation of the difference in relative cell content in the bacterium *Escherichia coli* Wyl. Raw data for Wyl is shown as red dots. Lack of difference is shown as a horizontal, black line and the 95% confidence of the lack of difference as a horizontal, dotted line. Insets in A and B, and raw data for relative drug content can be found in Figure S7.

## III. DICUSSION

Why do microbes form communities? It was recently (*5*) suggested that bacteria rarely work together based on the biological interactions between them. But fundamental physics could draw a different picture, and help answer the question. If, as my study suggests, two species growing together can tolerate better tetracycline than they would otherwise do isolated, it is not unreasonable to hypothesise that forming such communities can help diminish the effect of toxic molecules in nature. For example, to grow closer to an antibiotic-producing microorganism and benefit from the carbon available or tolerate chemotherapies. Or, indeed, taking over a particular niche in case of drug-tolerant microbes without producing specialised molecules such as bacteriocins (*39*). All as a byproduct of a physical law.

It should be stressed that my theory is not specific to *Escherichia coli* and tetracycline, so I anticipate that other species and molecules obey the same principle described here. An example can be found in the interactions between tumours and bacteria, and the resulting tolerance to chemotherapies (*8*); or between microbial pathogens (*6*)—albeit reliably predicting the outcome is still challenging (*5*). Now, there are limitations in the model some of which are imposed by experimental conditions. For simplicity, the Fick’s term in equations 1b and c has no spatial derivative as it would be expected. However, during the experiment I shoot the cultures to homogenise drug the distribution of tetracycline and nutriends. Here the aforementioned derivatived can be approximated the difference in drug molecules between environment and cytoplasm. The model also assumes reliance on a single carbon source. In reality, however, I used two: Glucose as the main carbon source, and casamino acids. The latter can indeed be used as carbon source when glucose is scarce (*40*). The concentration I used (0.1% w/v) was low enough to enable the growth of the microorganisms without showing diauxic growth dynamics typical from cultures that are sustained on multiple carbon sources (*41*). While speciation might indeed occur over time to avoid competition (*42*), is it really important to my conclusions?

The model that I present here is, like all models, wrong is some ways but also useful in others. The goal of my study is to demonstrate that growing alongside other microorganisms can predictably change the efficacy of a drug used against one—or against all—of them. It is, from the model standpoint, indifferent which carbon type is used to grow so long the relative frequency of the microbes surrounding the target is sufficiently high. Figure 1 shows the changes in IC_90_ are consistent regardless of the differences in growth parameters, akin to those existing between microorganisms using different carbon sources (*43*) or metabolic efficiencies (*19*). Moreover, the model and experiment use two species mixed in the same proportion for the purpose of pinpointing the physical mechanism. In nature, communities are more complex and this phenomenon likely to be more nuanced, if not even stronger, based on the ratio between the number of cells of the target species and its neighbours’.

My contribution with this model is to demonstrate that, contratry to current belief, the change in drug efficacy derived from the growth of other microorganisms is not arbitrary. There is, at least, one simple physical mechanism behind the changes in efficacy and, therefore, the changes can be qualitatively predicted—a necessary step if competition is to be harnessed (*5*) and used either in the clinic or treatment of pollutants in the soil.

## IV. METHODS

### Media and Strains

The strains of *Escherichia coli* GB(c) and Wyl were a gift from Remy Chait and Roy Kishony, and non-labelled *Salmonella, typhimurium* SL1344 a gift from Markus Arnoldini and Martia Ackermann. Experiments were conducted in M9 minimal media supplemented with 0.4% glucose (Fisher Scientific #G/0500/53) and 0.1% casamino acids (Duchefa #C1301.0250), supplemented with tetracycline. M9 minimum media (50X) was made by mixing equal proportions (vol/vol) of two parts, part A and part B, and diluted accordingly to 1X. Part A (50X) contains 350 g/L of K_2_HPO_4_(Sigma #P3786) and 100g/L of KH_2_HPO_4_(Sigma #P9791); whereas part B (50X) contains 29.4g/L of trisodium citrate (Sigma #S1804), 50g/L of (NH_4_)_2_SO_4_(Sigma #A4418), and 10.45g/L of MgSO_4_(Fisher Scientific #M/1050/53). I made tetracycline stock solutions from powder stock (Duchefa #0150.0025) at 5mg/mL in deionised water. Subsequent dilutions were made from this stock and kept at 4°C.

### Sensitivity assay

Using a 96-pin replicator, I inoculated a 96-well mifrotitre plate, containing 150μL of media supplemented with 0−0.5 μg/mL of tetracycline (for *E. coli* Wyl and *S*.*typhimurium*) or 0−15 μg/mL (for *E. coli* GB(c)), with 1 μL from an – overnight culture of each strain (approx. 2·10^8^ cells) to measure drug sensitivity in pure cultures. For sensitivity assays of Wyl in mixed culture conditions I inoculated the microtitre plate, containing 150 μg/mL of media supplemented with 0−0.5 μg/mL of tetracycline, with equal number of cells from two overnight cultures from Wyl and *S. typhimurium* using 8 technical replicates for each drug concentration. For the competition between Wyl and GB(c), however, I used 0−0.2 μg/mL to measure drug efficacy more reliably given the increased drug efficacy observed against Wyl. Figure S8 illustrates that mixing 1 μL of each species, therefore doubling the inoculum size with respect to pure culture conditions, or mixing 0.5 μL to maintain the same inoculum size has no impact on the sensitivity assays consistently with the non-linear relationship reported between inoculum size and drug efficacy (*31, 32*).

I incubated the microtitre plate at 30°C in a commercial spectrophotometer, shaking at 750rpm to ensure uniform mixing, and measured the optical density of each well at 600nm (OD_600_), yellow florescence for Wyl (YFP excitation at 505nm, emission at 540nm), and cyan fluorescence for GB(c) (CFP at 430nm/480nm) every 20min for 24h. I defined the minimum inhibitory concentration as the tetracycline concentration able to inhibit 90% of the growth observed in the absence of antibiotic after the 24h incubation period.

### Culture readings

Fluorescence protein genes were constitutively expressed with an approximately constant fluorescence to optical density ratio (Figure S9). The number of colony forming units (CFU) is positively correlated with optical density measured at 600nm (*OD*_600_) (Figure S10). Thus, I normalised fluorescence readings with respect to optical density readings, using the ratio optical density to fluorescence that I in pure culture conditions, to track the relative abundance of Wyl in mixed culture conditions. This metric is referred to as ‘Estimated optical density’ or ‘eOD’ in the main text. Time series data set were blank corrected prior to calculating the minimum inhibitory concentration.

## Code availability

A non-parameterized, python3 implementation of equations 1a-d can be found at https://github.com/rc-reding/papers/tree/master/MicrobCommunities_ISMECOMMS_2022. Table 1 contains a list of parameter values used.

**Table 1.** Model parameters for equations 1a–d, 2 and 3.

| Parameter | Description | Value |
| --- | --- | --- |
| $\bar{\mu}_j$ | Maximal carbon uptake rate | 1.25 mg / OD / h |
| $K_j$ | Half-saturation constant | 0.5 mg / mL |
| $y_j$ | Biomass yield | 0.65 OD / mg |
| $d$ | Drug degradation rate | $10^{-4}$ / h |
| $\kappa_j$ | Affinity of drug A for species type $j$ | 0.1 mL / μg |
| $\varphi_j$ | Diffusion coefficient | 10 μL / OD / h |
| $\alpha$ | Hill coefficient | 2 (dimensionless) |
| $A_0$ | Initial drug concentration | 2 μg / mL |
| $C_0$ | Initial carbon concentration | 2 mg / mL |
| $S_1(0)$ | Inoculum size for species $S_1$ | $10^{-3}$ OD |
| $S_2(0)$ | Inoculum size for species $S_2$ | $10^{-3}$ OD |

## Acknowledgements

The author thanks Robert Beardmore from the University of Exeter for laboratory support in the early stages of this study.

