## Supplementary material for "Predicting the re-distribution of antibiotic molecules caused by inter-species interactions in microbial communities"

### V. Supplementary Text

**Sensitivity modulation by species $S_2$ is independent of resistance mechanism.** In the main text, I defined whether species $S_2$ is sensitive to drug $A$ based on the numerical value of $\kappa_j$ in equation 3—which denotes the affinity of drug $A$ for its target. This is equivalent to mutations, say, in the bacterial ribosome that reduce the affinity of ribosome-binding antibiotics for their target (*1*). But other resistance mechanisms such, as efflux pumps, do not act on the binding affinity of a drug for its target. Efflux pumps can have broad specificity (*2*), thus, able to protect against multiple drugs simultaneously (*3, 4*).

To explore whether the predictions in the main text are robust to different mutation types, I redefined equations 1b and c to accommodate efflux pumps as resistance mechanism (*5*):

$$\dot{S}_j = \overbrace{G_j(C)S_j}^{\text{Growth}} \cdot \overbrace{I_j(A)}^{\text{Inhibition}}, \tag{4a}$$

$$\dot{A}_j = \overbrace{-dA_j}^{\text{Decay}} + \overbrace{\left(\varphi_j(A_e - A_j) - \frac{\upsilon^* p_j}{k_m^* + p_j}\right)S_j}^{\text{Fick's Diffusion with Efflux}}, \tag{4b}$$

$$\dot{A}_e = -dA_e - \sum_{j=1}^{n}\left(\varphi_j(A_e - A_j) - \frac{\upsilon^* p_j}{k_m^* + p_j}\right)S_j, \tag{4c}$$

$$\dot{C} = -\sum_{j=1}^{n} \overbrace{U_j(C)S_j}^{\text{C-Uptake}}, \tag{4d}$$

Here $\upsilon^*$ represents the maximal efflux rate; $k_m^*$ the half-saturation constant associated where the affinity of the pump for its substrate, $A$, is given by $1/k_m^*$; and $0 \le p_j \le 1$ is the expression level of $j-1$ copies of the efflux pump gene based on the limited abundance of DNA polymerase transcription complex (*6*). The abundance of this pump depends on the number of genes $j-1$ encoding efflux pump. The parameter $p_j$ is monotonically increasing and bounded in $j$, controlled by a dimensionless constant $\gamma$ in the Michaelis-Menten function $p_j = (j-1)/(1+\gamma(j-1))$ and $p_j/(k_m^* + p_j)$ the probability that a given drug molecule is bound to the pump. Thus, species $S_1$ does not express any efflux pump (no copies, as $j-1=0$) whereas species $S_2$ does indeed express the efflux pump. The remaining parameters are described in the main text with $\kappa_1 = \kappa_2$.

As Figure S11 illustrates, efflux pumps do not change the effect that competing genotypes have on species $S_1$. However, this resistance mechanism does increase even further the relative abundance of drug $A$ in species $S_1$. Drug $A$ also diffuses into competing species $S_2$ until it reaches equilibrium, and will diffuse back into the environment as the abundance of $A$ declines, effectively re-exposing $S_1$ (Figure S6G–I). Active efflux exacerbates this effect by actively moving $A$ molecules from within $S_2$ back into the environment. The result, shown in Figures S11A–C, is further inhibition of $S_1$ with respect to the mechanism used in the main text, as noted by its lower $IC_{90}$.

But not all resistant mutants rely on efflux pumps, nor they all help increase drug efficacy. Another common mechanism of resistance is enzymatic degradation (*7*). To mimic this effect, I implemented $d$ as another parameter that is independent for each species ($d_j$) in equations 1b and c.

The value of this parameter in the most sensitive species is the same as that assigned for the decay due to chemical stability of drug $A$ in the environment, so $d_e = d_1$. However, the competing species $S_2$ has $d_2 = 10,000 \times d_1$ to illustrate that, when inside this species, drug $A$ degrades substantially faster due to active enzymatic degradation. As it is to expect, degrading $A$ leads to an overall reduction in the availability of $A$ for all species and, therefore, to an increase in the $IC_{90}$ (Figure S12). This is consistent with experimental data where the presence of degraders within communities with antibiotic-producing microbes, allows drug-sensitive bacteria to grow closer to these microorganisms (*8*).

### VI. Supplementary Figures

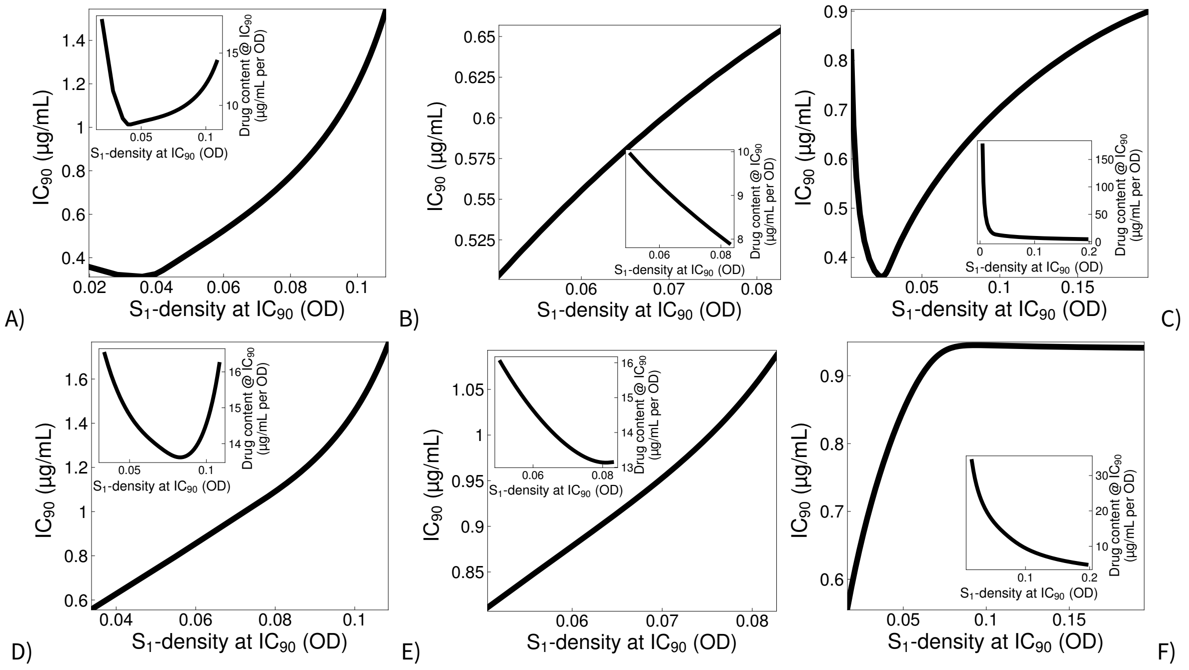


**Figure S1. Relationship between drug efficacy and $S_1$-cell density.** Variation in $IC_{90}$ (y-axis) as a function of cell density of species $S_1$ at its $IC_{90}$ (x-axis) when the incubation period is 24h (A–C) or 72h (D–F). The change in density is driven by higher maximal uptake rates ($\bar{\mu}$, A and D), carbon affinity ($k$, B and E), and biomass yield ($y$, C and F). In most cases there is a direct and non-linear correlation between both variables. The inset in each plot represents the drug content per $S_1$-cell at their $IC_{90}$ as a function of their density at the $IC_{90}$.

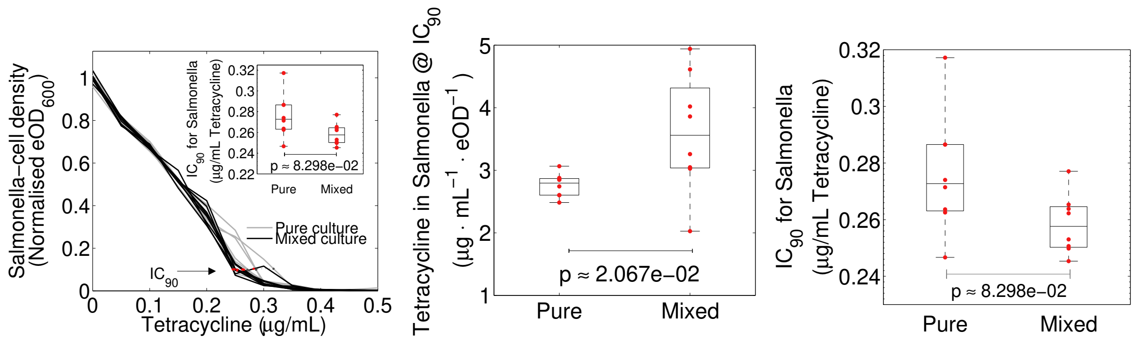


**Figure S2. Drug efficacy data for *Salmonella typhimurium*. Left)** Change in normalised density of *Salmonella typhimurium* as a function of tetracycline concentration, when Wyl grows in mixed culture with tetracycline-sensitive *Escherichia coli* Wyl. The change in density of *S. typhimurium* growing in mixed culture is shown in black, with grey showing the change in density in pure culture. The $IC_{90}$ in each condition is shown as dots, red for mixed culture conditions and dark grey for pure culture, connected by a dotted line. Non-parametric, Mann-Whitney U-test between $IC_{90}$s is shown in the inset. Raw data is shown as dots, whereas the boxes represent median (centre of the box), 25th, and 75th percentile of the dataset. The whiskers show the most extreme data points that are not outliers. Similar box plots show the relative content of tetracycline in *S. typhimurium* at its $IC_{90}$ (**centre**), and changes in $IC_{90}$ berween pure and mixed culture conditions (**right**)

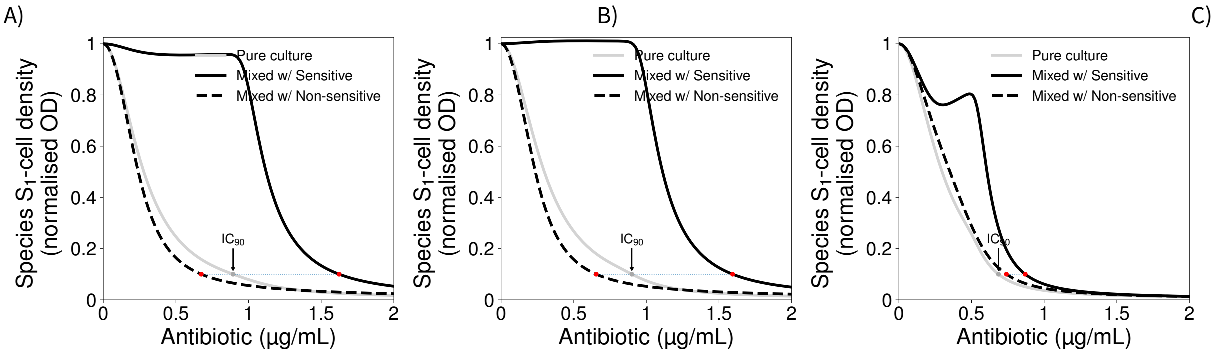


**Figure S3. Drug efficacy against species $S_1$ for different parameter values in species $S_2$.** Change in normalised density of species $S_1$, complementing Figure 1 in the main text, when A) the parameter for carbon uptake rate of species $S_2$ is 1.75 mg/OD/h, and when that for biomass yield is B) 0.35 OD/mg, and C) 0.75 OD/mg.

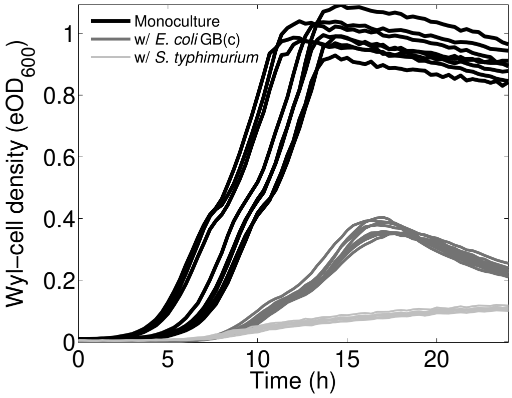


**Figure S4. Growth of *E. coli* Wyl without tetracycline.** Change cell density of tetracycline-sensitive construct Wyl, aka species $S_1$, complementing Figure 2 in the main text, when it is grown in pure culture (black), mixed with tetracycline-resistant construct of *E. coli* GB(c) (dark grey), and mixed with tetracycline-sensitive *S. typhimurium* (light grey).

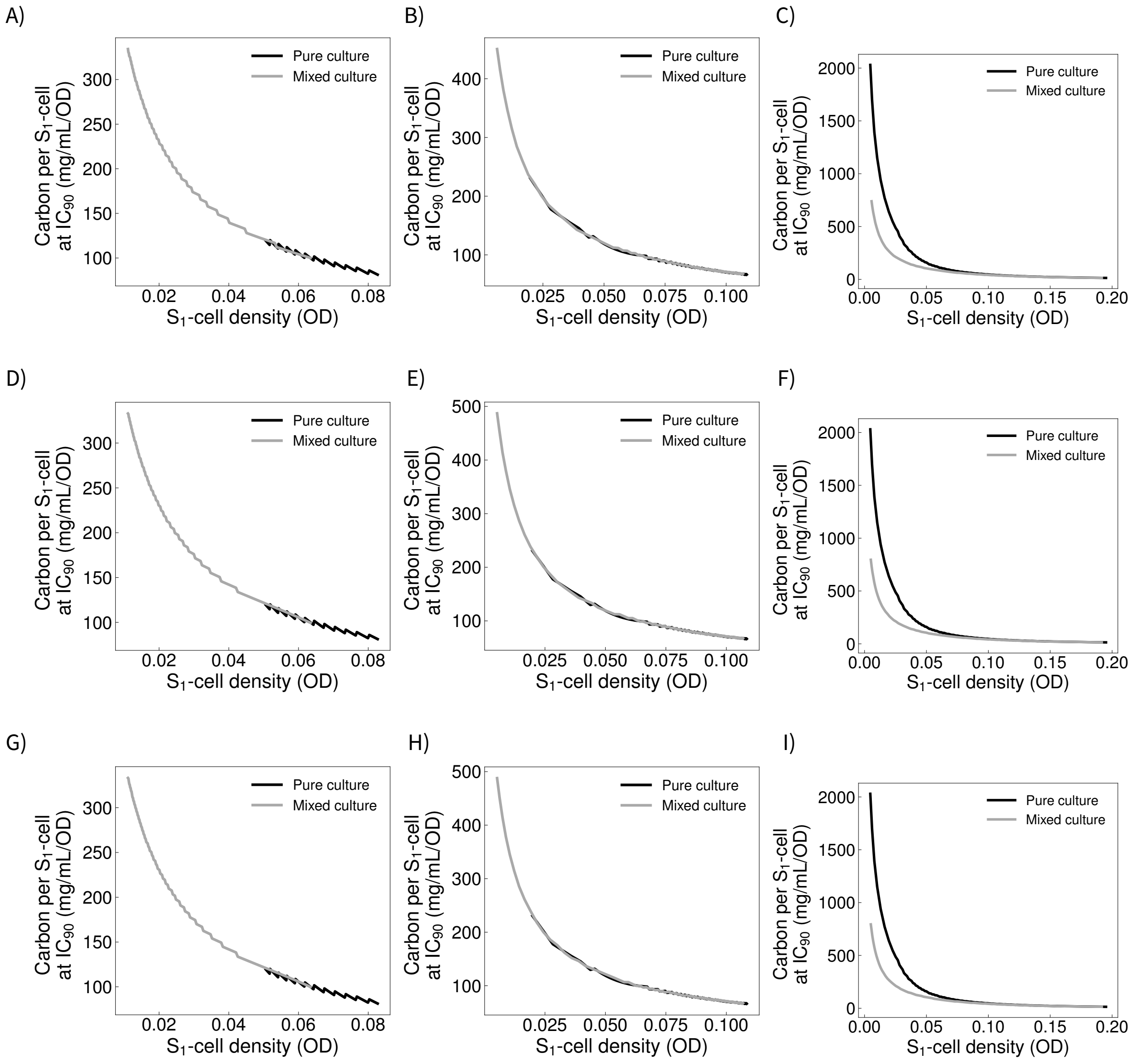


**Figure S5. Relative carbon molecules in species $S_1$ at the $IC_{90}$.** The plots represent the theoretical change in carbon molecules per $S_1$-cell, on the y-axis, as a function of cell density on the x-axis. The change predicted in the absence of neighbours is shown in back, whereas that observed growing alongside species $S_2$ is shown in grey. Figures A–C show the theoretical change in carbon molecules when neighbouring species $S_2$ is drug-sensitive, D–F the change when $S_2$ is drug-tolerant, and G–I the change when species $S_2$ is able to remove drug A from the cytoplasm through efflux pumps. Figures A, D, and G show the relative content of carbon for different carbon affinities (each OD point corresponds to a different parameter value); Figures B, E, and H for different maximal carbon uptake rates; and Figures C, F, and I for different biomass yield.

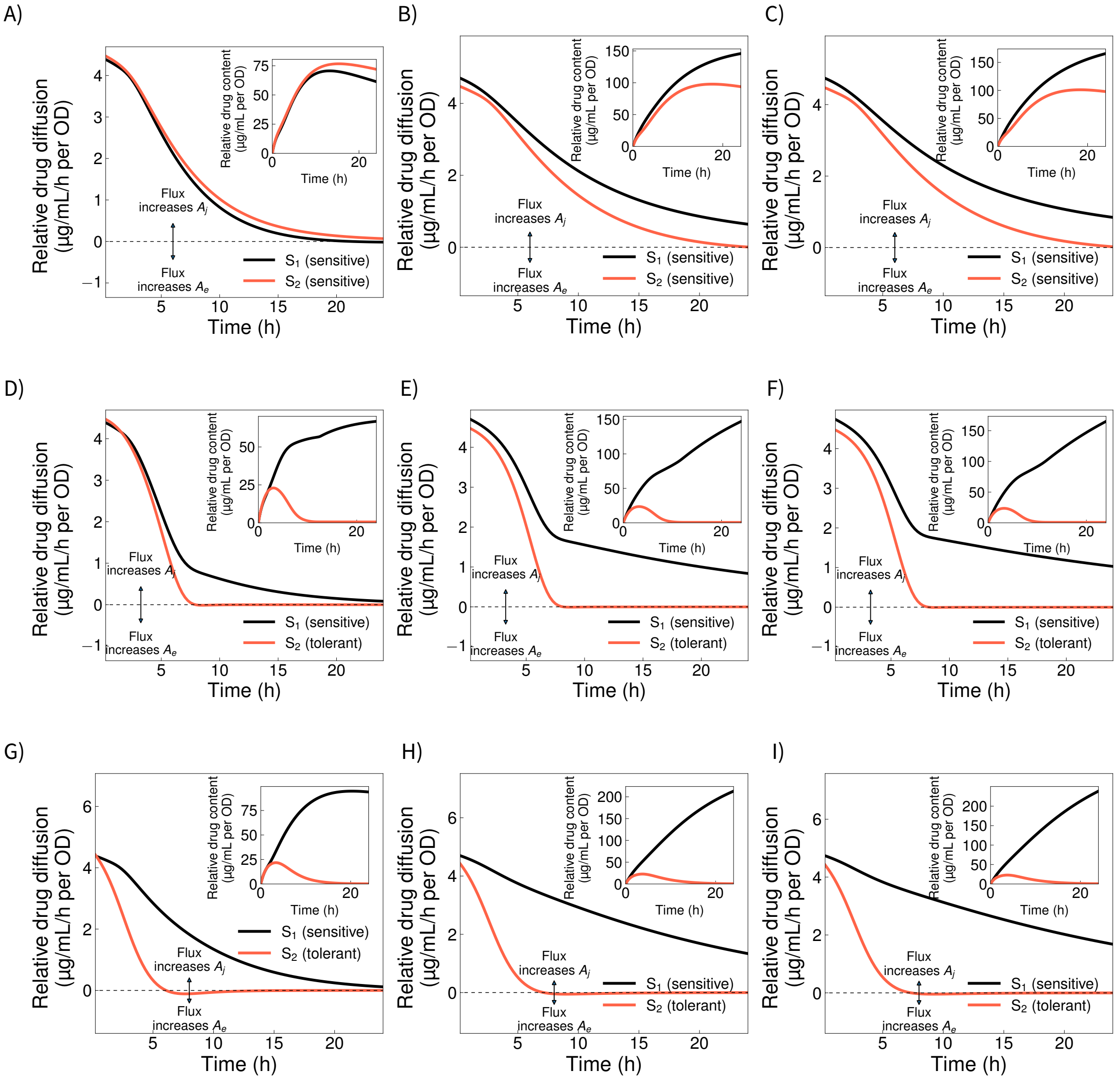


**Figure S6. Drug diffusion profiles for species $S_1$ and $S_2$ in mixed growth conditions.** The main plot shows the change in diffusion of drug $A$ over time, for competing species $S_1$ and $S_2$, as modelled by equations 1b (A–F) and 4b (G–I). The dashed line represents the point where both cytoplasm and environment have the same amount of $A$ and therefore in equilibrium ($A_j' = 0$). Thus, diffusion values above and below zero note that $A$ increases or decreases within each species, respectively. The inset shows the change in relative $A$ through time within each species.

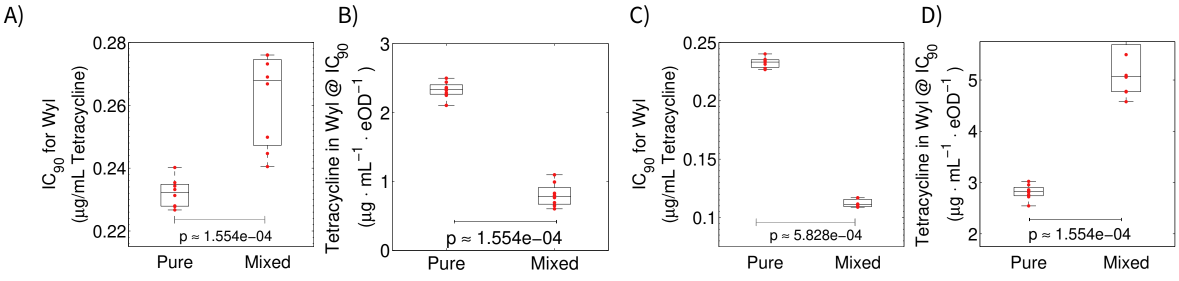


**Figure S7. Box plots for tetracycline efficacy against *Escherichia coli*.** The boxes represent median (centre of the box), 25th, and 75th percentile of the dataset for $IC_{90}$ and drug content in *Escherichia coli* Wyl growing alongside drug-sensitive *Salmonella typhimurium* (A and B), and drug-resistant *Escherichia coli* GB(c). The whiskers show the most extreme data points that are not outliers. The $p$ values shown correspond to non-parametric Mann-Whitney U-tests to determine significance of the difference in datasets between pure and mixed culture conditions.

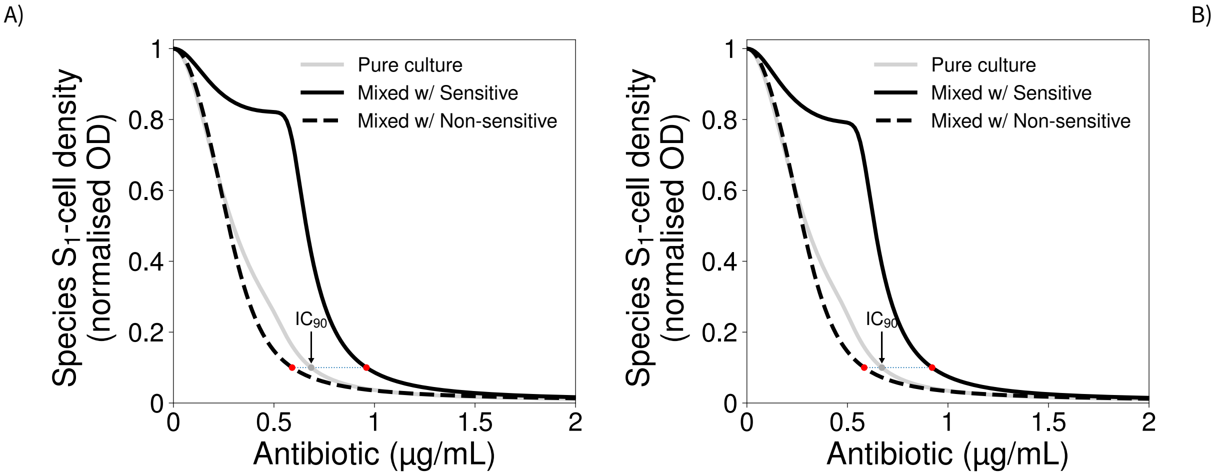


**Figure S8. Effect on drug efficacy of a 2-fold difference in inoculum size.** Theoretical change in $S_1$-cell density with increasing antibiotic concentration in pure (grey) and mixed (black) culture conditions with neighbours that have different drug sensitivity. The inoculum used for the mixed culture in **A)** is twice the size of that used in pure culture, by using the same sizes of $S_1$ and $S_2$ used in pure culture. The inoculum of each species is halved in **B)**, and therefore the inoculum size in pure and mixed cultures is identical.

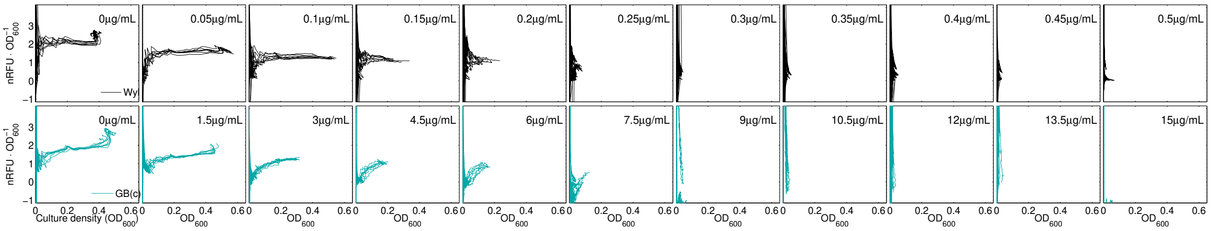


**Figure S9. Changes in relative fluorescence over time in both Wyl and GB(c) strains of *Escherichia coli*.** Raw change in florescence, per optical density units, measured every 20min for 24h for *E. coli* Wyl (black) and GB(c). Each column represents the data set for each tetracycline concentration used.

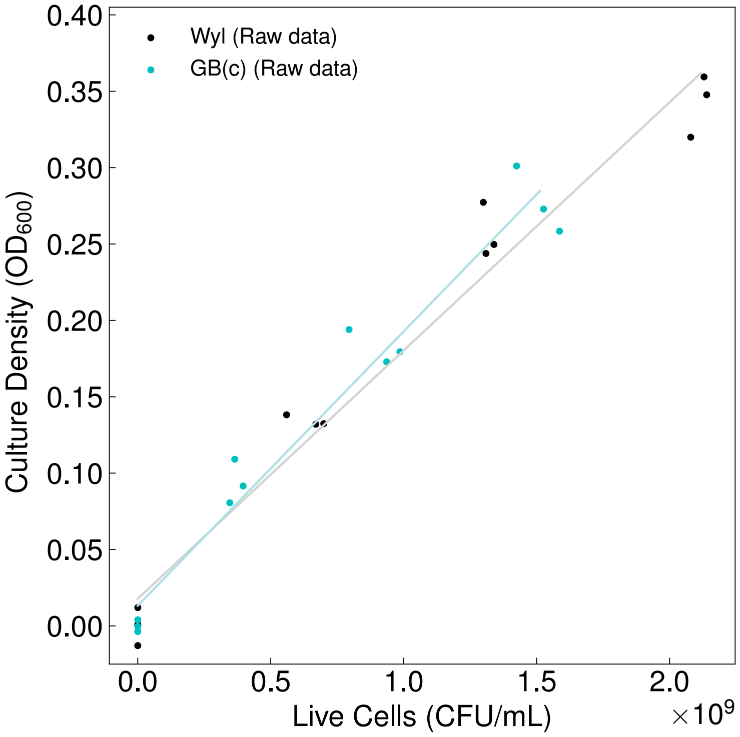


**Figure S10. Calibration curve to translate optical density data to number of *Escherichia coli* cells.** I fitted the linear model $a = bx + c$ to optical density and colony counting data (dots) to calculate the number of optical density units ($OD_{600}$) per cell. $a$ denotes the optical density readings measured at 600nm, $c$ the crossing point with the $y-$axis when $x = 0$, and $b$ the conversion factor between optical density and number of cells ($x$). I interpolating optical density readings to calculate the number of cells within a culture as $x = (a - c)/b$. For the strain S, $b = 1.62 \times 10^{-10}\ OD \cdot mL \cdot CFU^{-1}$ and $c = 1.78 \times 10^{-2}\ OD$, whereas for R $b = 1.79 \times 10^{-10}\ OD \cdot mL \cdot CFU^{-1}$ and $c = 1.33 \times 10^{-2}\ OD$.

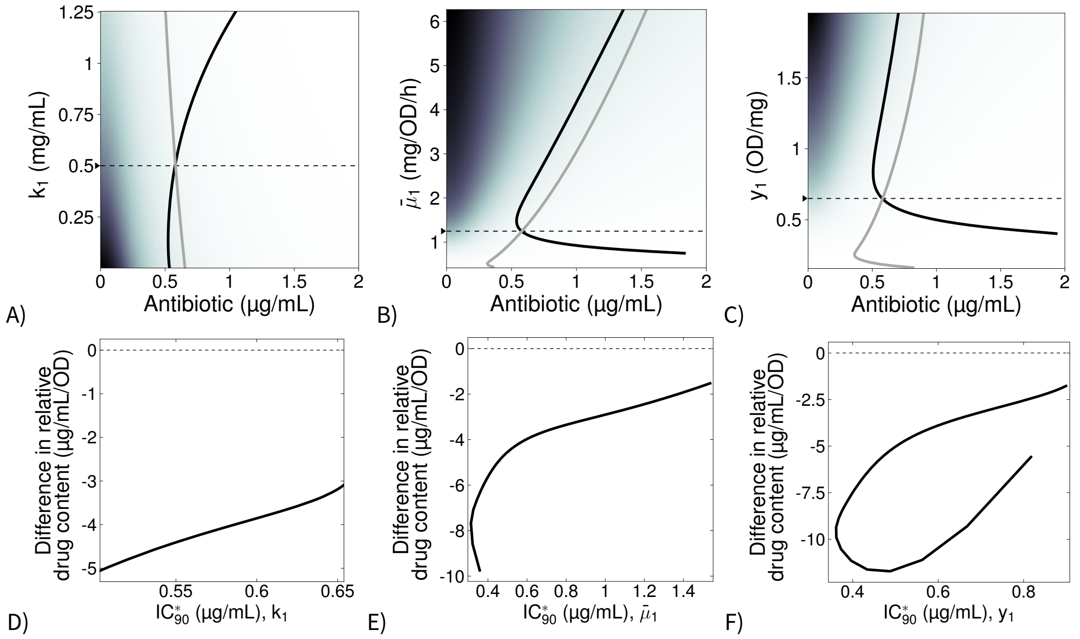


**Figure S11. Drug concentration in individuals from species $S_1$ in pure and mixed growth conditions when competing genotypes avoid drug-inhibition through efflux pumps. A–C)** $IC_{90}$, antibiotic concentration inhibiting 90% ($IC_{90}$) the growth predicted without drug, resulting with different parameters values for the half-saturation parameter $k_1$ (B), maximal carbon up-take $\bar{\mu}_1$ (C), or biomass yield $y_s$ (D) in equation 1 when species $S_2$ is drug-resistant through efflux pumps. The $IC_{90}$ for species $S_1$ growing as pure cultures is shown in grey, and growing in mixed culture with $S_2$ are shown in black. The parameter values for species $S_2$ were fixed at a value noted by a black arrow on the $y$-axis, followed by a dotted black line. **D–F)** Theoretical difference in relative drug content—antibiotic molecules per cell–of $S_1$ between pure culture conditions, and mixed culture with *drug-insensitive* $S_2$. D), E) and F) illustrate the prediction when changing the parameter $k$, $\bar{\mu}$, and $y$, respectively. The difference is positive (>0) when the relative content of antibiotic is higher in pure culture conditions, whereas is negative (<0) when the content is higher in mixed culture conditions. Lack of difference is represented by a horizontal, dotted line.

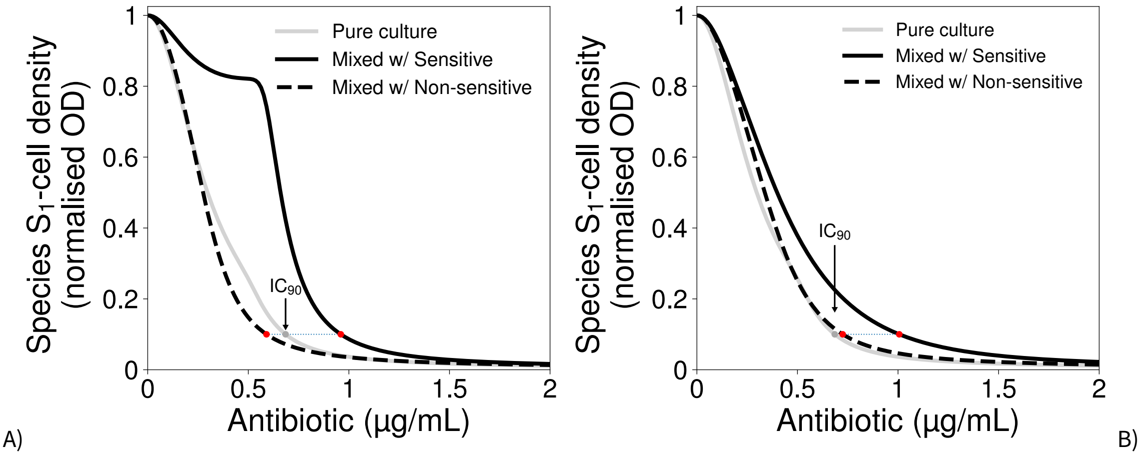


**Figure S12. Effect of competitors that can degrade drug A on the $IC_{90}$ of species $S_1$.** Theoretical change in $S_1$-cell density with increasing drug concentration in pure (grey) and mixed (black) culture conditions, with neighbours that have different drug sensitivity. The $IC_{90}$ in each condition is shown as dots, red for mixed culture conditions and dark grey for pure culture, connected by a dotted line. The plot in A) represents the case where both competitors have identical drug decay $d$ but different carbon uptake rate ($\bar{\mu_j}$). The plot in B) represents the case where $d_2 = 10,000\times d_1$ to emulate the competing, non-sensitive species being able to actively degrade the drug.
